# Responses of root architectural and anatomical traits to low nitrogen stress in rice

**DOI:** 10.1101/2024.08.28.610142

**Authors:** Tian Tian, Jonathan P. Lynch, Kathleen M. Brown

**Author notes:** National Academy of Forestry and Grassland Administration, Beijing 102600 China (Tian Tian).

## Abstract

Improving nitrogen use efficiency in rice would provide economic and environmental benefits, but little is known about root morphological and anatomical responses to low nitrogen. In this study, two sets of rice genotypes, one set from the RDP1 panel, and one set of recombinant inbred lines, were used to characterize responses to gradual nitrogen depletion by plant uptake and movement of nitrogen to deeper soil strata as a result of leaching, so that more nitrogen was available at depth in aerobic soil mesocosms in a greenhouse. There was significant genetic variation in shoot biomass reductions in response to low nitrogen. The root to shoot biomass ratio was increased by low nitrogen in both sets of genotypes. Relative investment in nodal root number was accentuated with low nitrogen, and shoot biomass was correlated with numbers and lengths of nodal and large lateral roots. There was genetic variation for nodal root number and length in both sets of genotypes. Anatomical responses to low nitrogen were assessed in nodal roots of the RILs, where root cross-sectional area, stele area, and metaxylem vessel number were reduced by low nitrogen, and root diameter was reduced in the RDP1 genotypes. There were significant interactions of nitrogen with genotype for stele area and percent aerenchyma in the RILs. Genetic variation for low nitrogen responses may be useful for selection of rice lines with greater nitrogen acquisition under nitrogen-leaching conditions.

## Background

Rice (*Oryza sativa* L.) a model monocotyledon species, is one of the most important crops for more than half of the global population. It is particularly important as a staple food for the 560 million poor people in Asia and consumption continues to rise with population growth (GRSP: Global Rice Science Partnership 2013). Rice production requires high water availability and fertility (Kekulandara et al. 2018). Nitrogen is the most abundant nutrient in plant tissues, and N fertilizer application is a major economic cost for rice farmers, especially in developing countries (Vinod and Heuer 2012). Plants take up and assimilate both nitrate and ammonium, but nitrate is the predominant form in most agricultural soils.

Crop plants use only about 30% of the nitrogen applied to the soil (Raun and Johnson 1999; Sylvester-Bradley and Kindred 2009). Thus, more than 60% of the soil nitrogen is lost through a combination of leaching, surface run-off, denitrification, volatilization, and microbial consumption (Robertson and Vitousek 2009). In rice, ammonia volatilization is the major path of N loss, and N use is estimated to be 30-50% less efficient than what is achievable with good management (GRSP: Global Rice Science Partnership 2013). In upland rice production, nitrogen loss to leaching is expected to be greater than in paddy systems, and similar that of other crops, i.e. rainfall would carry nitrogen below the root foraging zone when plant uptake lags behind movement of nitrogen through the soil (Lynch 2013).

Root system architecture is a primary determinant of soil exploration, and thereby the efficiency and effectiveness with which a plant captures water and nutrient resources (Lynch 1995, 2013; Rebouillat et al. 2009; Gowda et al. 2011). Root systems should be efficient, i.e. use internal resources effectively to acquire soil resources, since they depend on the shoot for fixed carbon and must acquire limiting resources from different soil domains. For example, phosphorus (P) is usually more available in the shallower soil layers, while many drought scenarios result in more moisture availability at depth. Nitrogen is initially most available in shallow soil following fertilizer application, but rain and irrigation causes leaching of N deeper into the soil. The *steep, cheap and deep* ideotype has been proposed for efficient acquisition of deeper soil resources including nitrogen and water by maize crops (Lynch 2013). In maize, fewer nodal (crown) roots were associated with deeper rooting, better nitrogen capture, and better yield under low N stress (Gao and Lynch 2016). However, a greater number of crown roots was associated with shallower rooting and better P capture under low P (Sun et al. 2018). Similarly, less lateral root branching density increased axial elongation, root depth, and grain yield under low nitrogen (Zhan and Lynch 2015) and drought stress (Zhan and Lynch 2015), while greater lateral root branching density was associated with better performance under low P (Jia et al. 2018). In maize, root cortical aerenchyma formation reduces respiratory costs per unit root length, making root length “cheaper”, and this was associated with reduced specific root respiration, greater rooting depth, and better yield under low nitrogen stress (Saengwilai et al. 2014a), drought (Zhu et al. 2010; Chimungu et al. 2015), and low P stress (Galindo-Castañeda et al. 2018). Maize genotypes that increased their crown root branching angles under low N stress showed greater root distribution at depth and better performance at two field sites (Trachsel et al. 2013).

Like other cereals, rice productivity is limited by nitrogen stress, and it has been suggested that root architectural plasticity, i.e. responses to nitrogen availability, could be an important target for improving nitrogen use efficiency in rice (Ogawa et al. 2014). The rice root system can be divided into different classes: seminal roots, embryonic nodal roots, postembryonic nodal roots, large lateral roots and small lateral roots (Rebouillat et al. 2009). Five embryonic nodal roots emerge from the coleoptile and subsequent postembryonic nodal roots emerge from the nodes at the base of the main stem and tillers (Coudert et al. 2010). Functionally, nodal roots constitute a framework for the whole root system. When nodal roots exceed a certain length, lateral roots are initiated from the root pericycle and emerge through the cortex and epidermal layers. Lateral roots, which comprise the greatest proportion of the root system in total length and number, are responsible for most water and nutrient absorption (Kirk 2003). Lateral roots can be classified into two main types, small (S-type) lateral roots with determinate growth, lacking gravitropism and secondary lateral roots, and large (L-type) lateral roots with indeterminate growth, positive gravitropism, and secondary branching (Rebouillat et al. 2009). This suggests different functions for these two types of lateral roots. Rice nodal root anatomy is characterized by a high proportion of aerenchyma within the root cortex, and development of apoplastic barriers at the Casparian band and the exodermis.

Although there has been substantial research on rice root phenotypic responses to drought stress (Kano et al. 2011; Uga et al. 2013; Sandhu et al. 2014; Kadam et al. 2017; Hazman and Brown 2018), less is known regarding rice root responses to low nitrogen. Most studies on nitrogen responses in rice roots have focused on uptake and transport mechanisms while only a few reports have described rice root architectural responses to low nitrogen availability, and most of these focus on complex traits such as total root length, root depth distribution, and branching as evaluated from the number of tips and crossings, e.g. (Ogawa et al. 2014; Ju et al. 2015). The responses of anatomical phenotypes to low nitrogen have likewise received little attention in rice. Exceptions include a report that root cortical aerenchyma increases with N deficiency (Abiko and Obara 2014), which is important for nitrification of ammonia to nitrate in flooded soils (Kirk 2003), and the reduced development of apoplastic barriers (lignin and suberin) with low N (Ranathunge et al. 2016).

The main objectives of this research are to evaluate the responses of root architectural and anatomical phenotypes to low nitrogen availability in rice under conditions that mimic nitrogen -leaching soils found in upland production systems. We hypothesize that (1) rice exhibits alterations in root architecture and anatomy in response to low nitrogen, and (2) there is genetic variation in root responses to low nitrogen.

## Material and methods

### Plant materials and experimental design

Experiments were conducted using rice genotypes from two different populations. Eight diverse tropical japonica genotypes from Rice Diversity Panel 1 (RDP1) (Zhao et al. 2011) were selected for RDP1 experiments (Table S1). Eight single-seed descent recombinant inbred lines (RILs 72, 93, 147, 159, 188, 303, 330, 339) from a cross of Azucena X IR64 (Ahmadi et al. 2005) and provided by Dr. Susan McCouch, Cornell University, were used in separate experiments. Each set of experiments used a randomized complete block design with two nitrogen levels (high and low nitrogen), eight genotypes and four replicates. Planting of replicates was staggered in time as a block effect.

The plants were grown in a temperature-controlled greenhouse in University Park, PA, USA (40°48′N, 77°51′W) with temperatures ranging from 23-31°C and daylength of 14 h provided by natural light and supplemental LED lamps providing 150 μM m^-2^ s^-1^ PAR. Rice seeds were surface sterilized with 5 % bleach and pre-germinated prior to planting. For germination, seeds were sown on moist paper towels soaked with 0.5 mM CaSO_4_ and incubated for 1 d at 28 °C. Seedlings were transplanted to each 1.2×0.15 m mesocosm containing a growth medium consisting of a mixture of 40% vermiculite, 40% sand (0.3-0.5mm), 5% perlite, and 15% sifted field topsoil (Hagerstown silt loam). The mesocosms were lined with transparent high-density polyethylene film to facilitate root extraction. Twenty-five liters of growth medium was used in each mesocosm to ensure the same bulk density of the medium. Two days before planting, the cylinders were irrigated with 3.5 L of a nutrient solution adjusted to pH 5.0. The nutrient solution for the high-N treatment consisted of (in μM): NO_3_ (1400), NH_4_ (1400), P (320), K (1020), Ca (1000), SO_4_ (2830), Mg (1700), B (57), Zn (0.15), Cu (0.15), Mo (0.28), and DTPA-Fe (3.75). For the low-N treatment, NO_3_ and NH_4_ were reduced to 14 μM each.

For the RDP1 experiment, seeds were transplanted on Feb 24, Mar 03, Mar 31, and Apr 05, 2017. For the RILs experiment, seeds were transplanted on Oct 20 and Oct 25, 2017 and Feb 28 and Mar 9, 2018. Three germinated seeds were planted in each mesocosm, and after 7 days they were thinned to one plant. For seedling establishment, 200 mL of nutrient solution with N were applied to the high-N mesocosms every 2 days for 8 days, then daily, via drip irrigation using a DI-16 Dosatron fertilizer injector (Dosatron International Inc, Dallas, TX, USA). In the low-N treatment, plants were supplied with 200 mL of nutrient solution without N, so that the only N available was what was already present in the topsoil. Leachate was collected weekly from 3 ports at 30, 60 and 90cm from the top of the mesocosm using a soil micro sampler 2.5 mm in diameter and 9 cm in length (Soilmoisture Equipment Corp, Santa Barbara, CA, USA). The concentration of nitrate in the solution was immediately determined using a Nitrate Meter (LAQUA Twin, 2305GL, Aurora, IL, USA). NO_3_ leaching was observed in the high N treatment, where the NO_3_ concentration increased gradually over time at the 60 and 90 cm sampling positions, while at 30 cm from the top, N concentration was maximal at 2 weeks and then declined, but always remained at concentrations greater than 50 mg L^-1^ (=1.09 M) (Supplemental Figure1). In the low nitrogen treatment, the NO_3_ concentration was greatest at 90 cm depth throughout the experiment, while the nitrogen concentrations at 30 and 60 cm were always less than about 50 mg L^-1^. By the end of the experiment, NO_3_ concentration at the upper sampling position was 0.32 M in the RDP1 experiment and 0.46 M for the RILs experiment (Supplemental Figure 1).

### Shoot and root growth and N content

Plants were harvested 5 weeks after transplanting. Shoot and root nitrogen content was evaluated in the RILs experiment. One day before harvest, the number of tillers and leaves and the leaf chlorophyll content (SPAD) was measured at the middle of the penultimate fully expanded leaf, and the average of three SPAD values for each plant is presented). At harvest, root systems in both experiments were excavated and washed as described below and preserved in 75% ethanol until the time of processing and analysis.

Shoots and roots were dried at 65°C for 72h prior to dry weight determination. The N concentration was determined by using an elemental analyzer (PerkinElmer Series II CHNS/O Analyzer 2400, Shelton, CT, USA).

### Root Architecture Measurements

At harvest, the polyethylene liner containing the root and growth medium was carefully removed from the mesocosm and placed on a screen. The maximum root length was recorded after carefully removing some medium from the bottom of the column before the plastic liner was split open. After opening the plastic liner and washing the growth medium from the root system, the number of nodal roots was counted. The nodal roots were categorized into thin and thick nodal root classes according to root diameter, approximately > 0.35 mm for thin nodal root and > 0.6 mm for thick nodal root, with minor adjustments based on visual determination of root class, lateral root branching density and color. Three thin and three thick nodal roots representing the range of nodal root length were collected from each plant for measurements of nodal root length, and the middle segments (50-81.5 cm for HN and 13-78 cm for low nitrogen, measured from the root base) of the three thin nodal roots were used for measurement of small (S-type) and large (L-type) lateral root length and lateral root branching density. Only thin nodal roots were harvested in the RDP1 experiment. Root samples were then preserved in 75% ethanol for root architecture analysis. Small lateral root branching density was defined as the density of small lateral roots borne on both nodal roots and on large lateral roots per unit length of the subtending axes, and large lateral root branching density was defined as the number of large lateral roots per unit length of nodal root. Thin nodal roots were divided into 20 cm segments starting from the root base to measure axial distribution of root traits. The nodal roots were scanned and analyzed using WinRhizo software (WinRhizo Pro; Régent Instruments, Québec, Canada), and roots were classified using diameter classes for small and large lateral roots of < 0.055 mm and 0.055– 0.55 mm for HN and < 0.045 mm and 0.045– 0.35 mm for low nitrogen, respectively, since lateral root diameters were typically less in low nitrogen. Total root length was estimated from the weights of the scanned samples by the formula: total root length = total root weight / scanned root weight × scanned root length.

### Root anatomy measurements

In the RILs experiment, three 2-cm root samples were collected at 20, 40 and 60 cm from the thin nodal root base, and at 20 cm from the base of thick nodal roots. We also collected large lateral root samples from each thin nodal root segment. The root segments were placed into white plastic histocaps and stored in 70% ethanol. The samples were dehydrated in 80%, 90% and 100% ethanol for 1 hour each, and then the samples in histocaps were critical point dried according to manufacturer instructions (Leica EM CPD300, Buffalo Grove, IL). The critical-point dried root samples were used for anatomical analysis by laser ablation tomography (Hall et al. 2019; Strock et al. 2019). Briefly, laser ablation tomography uses a laser beam (Avia 7000; 355 nm pulsed laser) to vaporize or sublimate the root samples on a stage that advances towards a camera (Canon T3i with a 5X MP-E 65 mm microlens), which records cross-sectional images during ablation. Three best images per sample were selected and analyzed with MIPAR software (Sosa et al. 2014) for quantitative analysis of root cross section area (RCSA), aerenchyma area (AA), percent aerenchyma (AA%), total stele area (TSA), mean metaxylem vessel area (MXA), and number of metaxylem vessels (MXV).

### Statistical analyses

Empirical data were analyzed using SPSS 20.0. Two-way ANOVA was used to analyze the effects of nitrogen treatment, genotype, and their interactions (N×G) on shoot and root morphological and anatomical traits for thin and thick nodal roots, and three-way ANOVA was used in experiments involving multiple root axial positions. Pearson correlation was used to analyze the correlation between root traits and shoot biomass. Multiple comparisons were made using Tukey’s honestly significance difference (HSD) test. In all tests, a p value less than 0.05 was considered statistically significant unless otherwise noted.

## Results

### Growth and nitrogen accumulation

Low nitrogen treatment was effective in producing nitrogen stress in both experiments, as shown by significantly reduced shoot growth, root dry weight, SPAD value, and shoot and root nitrogen content (Figures 1-2, Table 1, Table S2). Low nitrogen treatments reduced shoot biomass, tiller number and leaf number by 79%, 60% and 62%, respectively, in the RILs experiment (Table 1) and to a similar extent in the RDP1 experiment, where plants generally had less biomass (Table S2). Significant nitrogen treatment × genotype interactions were observed for shoot biomass and number of leaves and tillers in both experiments. Reductions in shoot biomass with low nitrogen ranged from 69% in RIL 339 to 84% in RILs 72 and 303 in the RIL experiment, with similar ranges of response in the RDP1 experiment (Figure 1). Since low nitrogen reduced shoot biomass to a greater extent than root dry weight, root to shoot ratio was increased significantly under low nitrogen (Table 1, Table S2). Although low nitrogen significantly reduced shoot and root N content (Table 1), shoot nitrogen content per unit root length or weight increased approximately four-fold under low nitrogen treatment compared with high nitrogen (Table S3).

**Figure 1.**
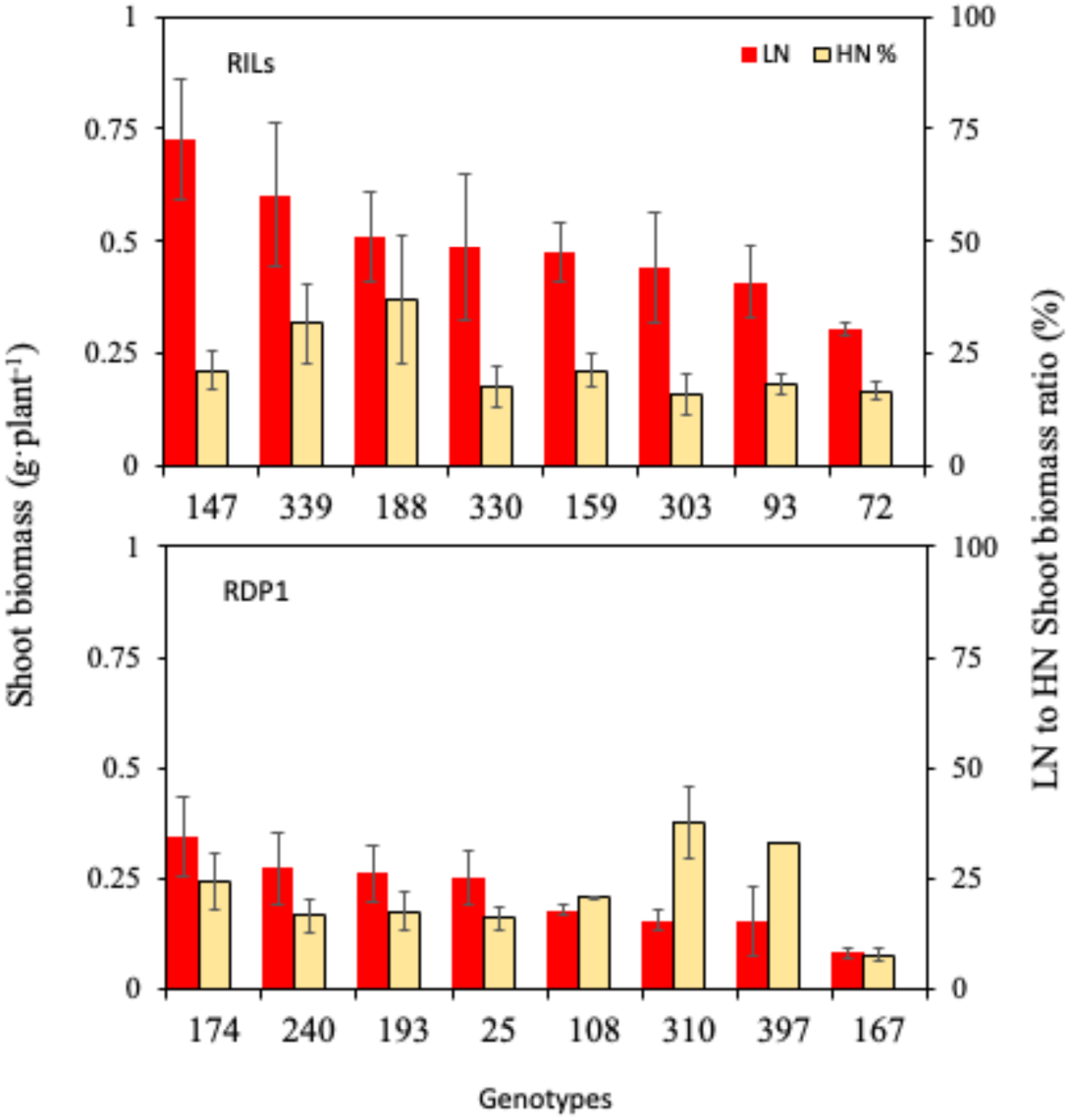
Shoot biomass with low N treatment and proportion of high N shoot biomass (LN as % HN) for each genotype in the RIL and RDP1 experiments. Values shown are means ± SE.

**Figure 2.**
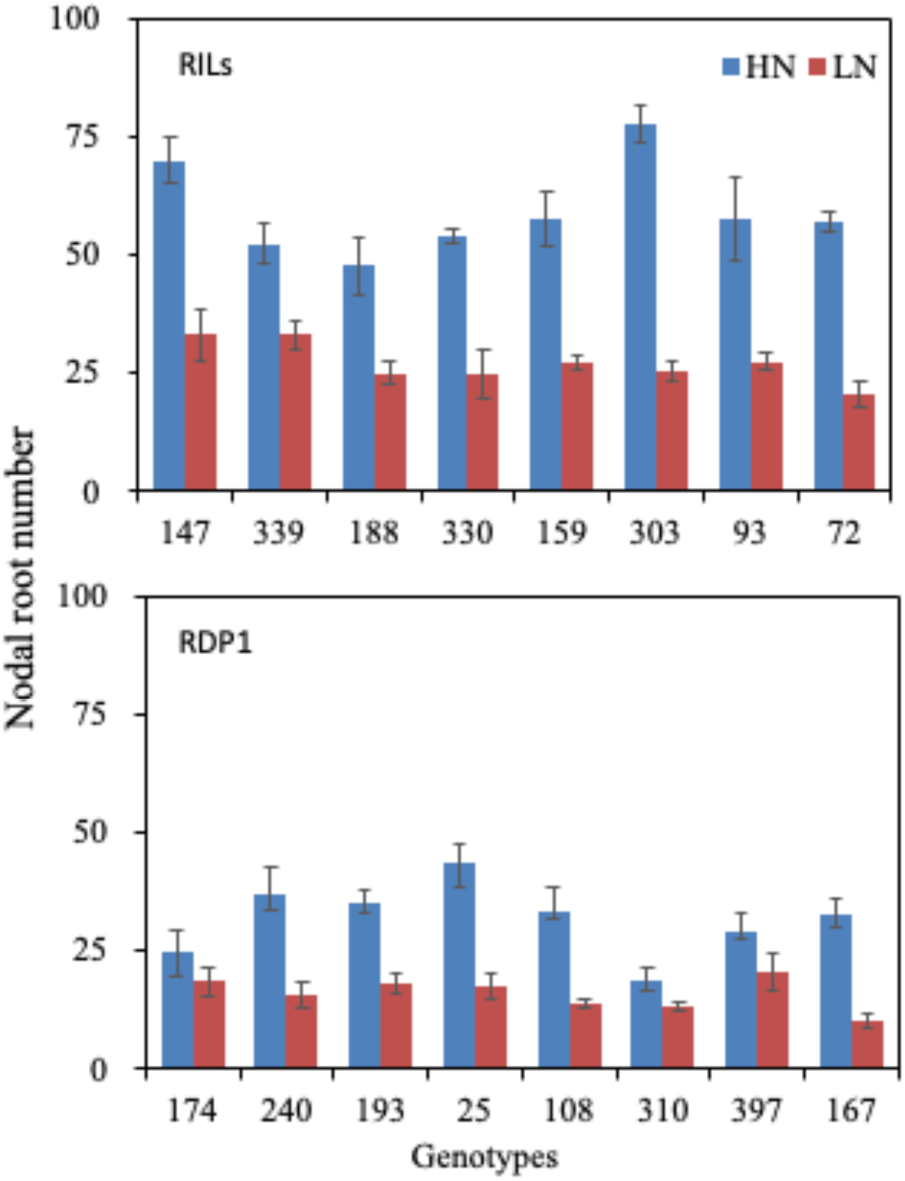
Effect of nitrogen treatment on nodal root number of each genotype in the RIL and RDP1 experiments. Values shown are means ± SE.

**Table 1.** The effects of nitrogen treatment and genotype on shoot and root morphological phenotypes of RILs.

|  | Shoot biomass (g·plant<sup>-1</sup>) |  | Plant height (cm) |  | Number of leaves (plant<sup>-1</sup>) |  | Number of tillers (plant<sup>-1</sup>) |  |
| --- | --- | --- | --- | --- | --- | --- | --- | --- |
|  | F | P | F | P | F | P | F | P |
| Treatment (N) | 349.084 | <0.001 | 83.604 | <0.001 | 293.879 | <0.001 | 277.204 | <0.001 |
| Genotype (G) | 4.852 | <0.001 | 3.813 | 0.002 | 10.249 | <0.001 | 5.788 | <0.001 |
| N*G | 2.966 | <0.001 | 1.487 | 0.194 | 3.499 | 0.004 | 2.527 | 0.027 |
| Treatments | Mean | SE | Mean | SE | Mean | SE | Mean | SE |
| HN | 2.398 a | 0.124 | 56.25 a | 1.234 | 35.9 a | 1.79 | 11.06 a | 0.469 |
| LN | 0.496 b | 0.041 | 44.00 b | 0.969 | 13.8 b | 0.83 | 4.41 b | 0.233 |
|  | Root dry weight (g·plant<sup>-1</sup>) |  | Root:shoot ratio |  | Maximum root length (cm) |  | SPAD |  |
|  | F | P | F | P | F | P | F | P |
| Treatment (N) | 140.197 | <0.001 | 11.567 | 0.001 | 5.313 | 0.026 | 193.728 | <0.001 |
| Genotype (G) | 7.028 | <0.001 | 2.503 | 0.028 | 1.607 | 0.156 | 4.156 | 0.001 |
| N*G | 3.533 | 0.004 | 0.914 | 0.504 | 0.912 | 0.505 | 1.612 | 0.155 |
| Treatments | Mean | SE | Mean | SE | Mean | SE | Mean | SE |
| HN | 1.129 a | 0.092 | 0.46 b | 0.02 | 73.6 a | 1.951 | 39.95 a | 0.626 |
| LN | 0.299 b | 0.034 | 0.58 a | 0.03 | 67.4 b | 1.994 | 28.90 b | 0.712 |
|  | Nodal root number (plant<sup>-1</sup>) |  | Nodal root length (cm) |  | Thin nodal root length (cm) |  | Small lateral root length per thin nodal root (cm) |  |
|  | F | P | F | P | F | P | F | P |
| Treatment (N) | 229.876 | <0.001 | 19.342 | <0.001 | 19.862 | <0.001 | 1.544 | 0.220 |
| Genotype (G) | 3.475 | 0.004 | 2.822 | 0.015 | 3.523 | 0.004 | 0.885 | 0.525 |
| N*G | 2.273 | 0.044 | 0.806 | 0.586 | 0.892 | 0.520 | 1.164 | 0.341 |
| Treatments | Mean | SE | Mean | SE | Mean | SE | Mean | SE |
| HN | 59.34 a | 2.30 | 62.7 a | 1.51 | 61.9 a | 1.50 | 460.7 a | 41.39 |
| LN | 26.41 b | 1.20 | 49.5 b | 2.89 | 52.3 b | 1.91 | 375.0 a | 55.45 |

|  | Large lateral root length per<br>thin nodal root (cm) |  | Thick nodal root length (cm) |  | Small lateral root length per thick<br>nodal root (cm) |  | Large lateral root length per<br>thick nodal root (cm) |  |
| --- | --- | --- | --- | --- | --- | --- | --- | --- |
|  | F | P | F | P | F | P | F | P |
| Treatment (N) | 2.749 | 0.104 | 2.220 | 0.143 | 0.433 | 0.514 | 3.662 | 0.062 |
| Genotype (G) | 1.430 | 0.216 | 0.910 | 0.507 | 0.427 | 0.880 | 0.824 | 0.572 |
| N*G | 0.633 | 0.727 | 0.353 | 0.925 | 1.509 | 0.187 | 1.139 | 0.355 |
| Treatments | Mean | SE | Mean | SE | Mean | SE | Mean | SE |
| HN | 702.7 a | 55.79 | 20.7 a | 1.38 | 22.8 a | 3.17 | 38.7 a | 9.40 |
| LN | 571.7 a | 56.39 | 17.9 a | 1.09 | 19.7 a | 3.34 | 17.0 a | 6.30 |

Analysis of variance (ANOVA), means and standard errors (SE) are shown for 8 genotypes with high nitrogen (1400 μM; HN) and low nitrogen (14 μM; LN) treatments.
Different letters indicate significant difference ( $p<0.05$ )

### Root architectural responses to low nitrogen

Nodal root number was significantly reduced by low nitrogen, and the extent of reduction varied among genotypes (e.g. from 44% in RIL 188 to 68% in RIL 303) (Table 1, Table S2, Figure 2). Thin nodal roots were longer than thick nodal roots in both high and low nitrogen, and thin nodal roots bore greater lengths of small and large lateral roots than thick nodal roots (Table 1). Thin, but not thick, nodal roots were significantly shorter under low nitrogen. Neither genotype nor N treatment affected lateral root length per nodal root (Table 1), or small or large lateral root branching density assessed over the entire nodal root sample (data not shown). However, lateral roots were not uniformly distributed along the nodal root axes (Table 2, Table S4, Figures 3-5). Large lateral root lengths and branching densities were typically greatest in the upper root segments in both high and low nitrogen (Table 2, Table S4, Figures 3-5). RDP1 genotypes, but not RILs, had significant nitrogen treatment X genotype interactions for large lateral root length and branching density (Table 2, Table S4).

**Figure 3.**
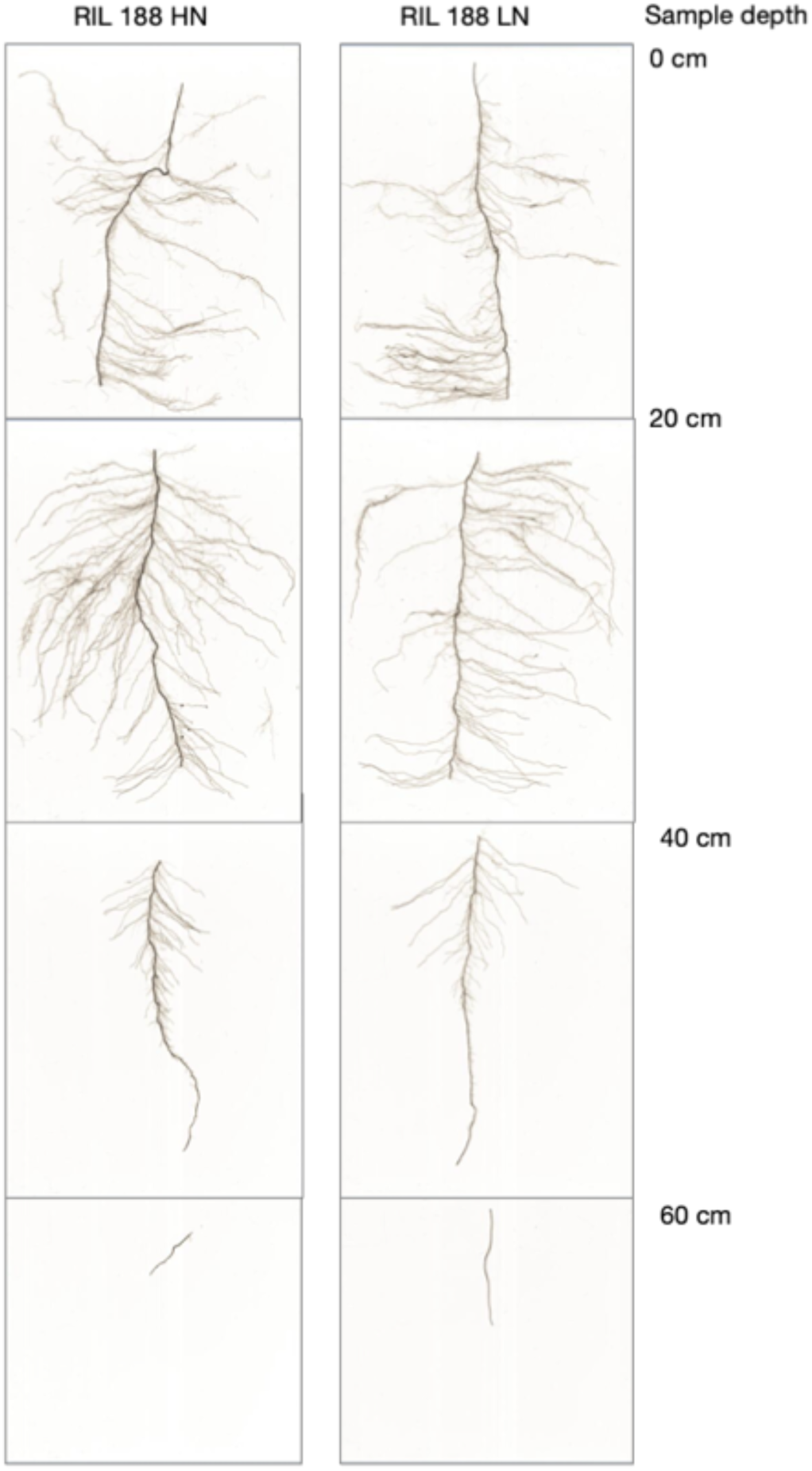
Typical distribution of lateral roots along nodal root axes of RIL188 grown under high and low nitrogen. Note the maintenance of deep rooting despite the reduction in overall plant biomass.

**Figure 4.**
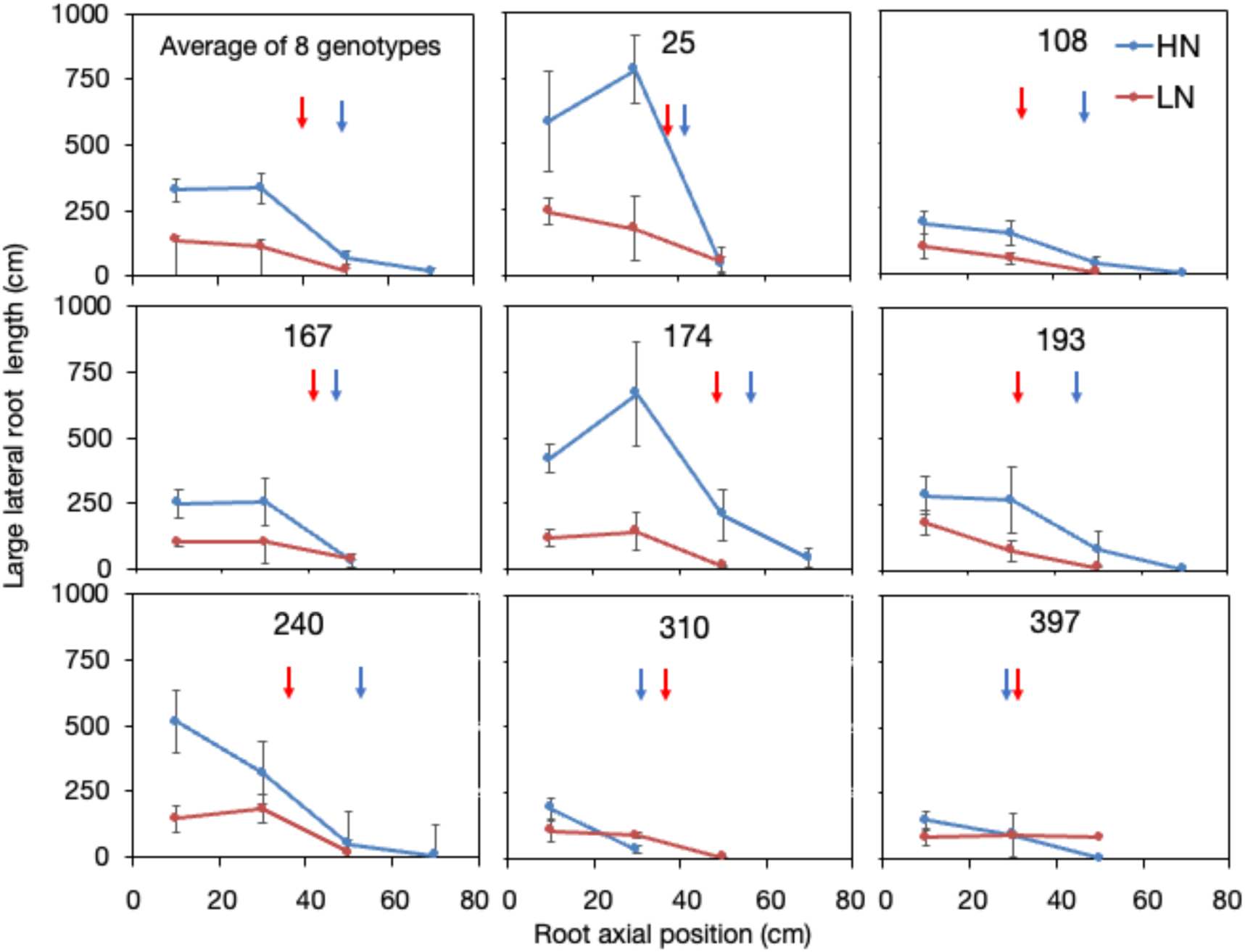
Effect of nitrogen treatment on distribution of large lateral root length along nodal root axes of each genotype in the RDP1 experiment. Values shown are means ± SE. Where no error bar is shown, there was only one nodal root sample at that depth. Arrows indicate mean nodal root length for high N (blue) and low N (red) treatments.

**Figure 5.**
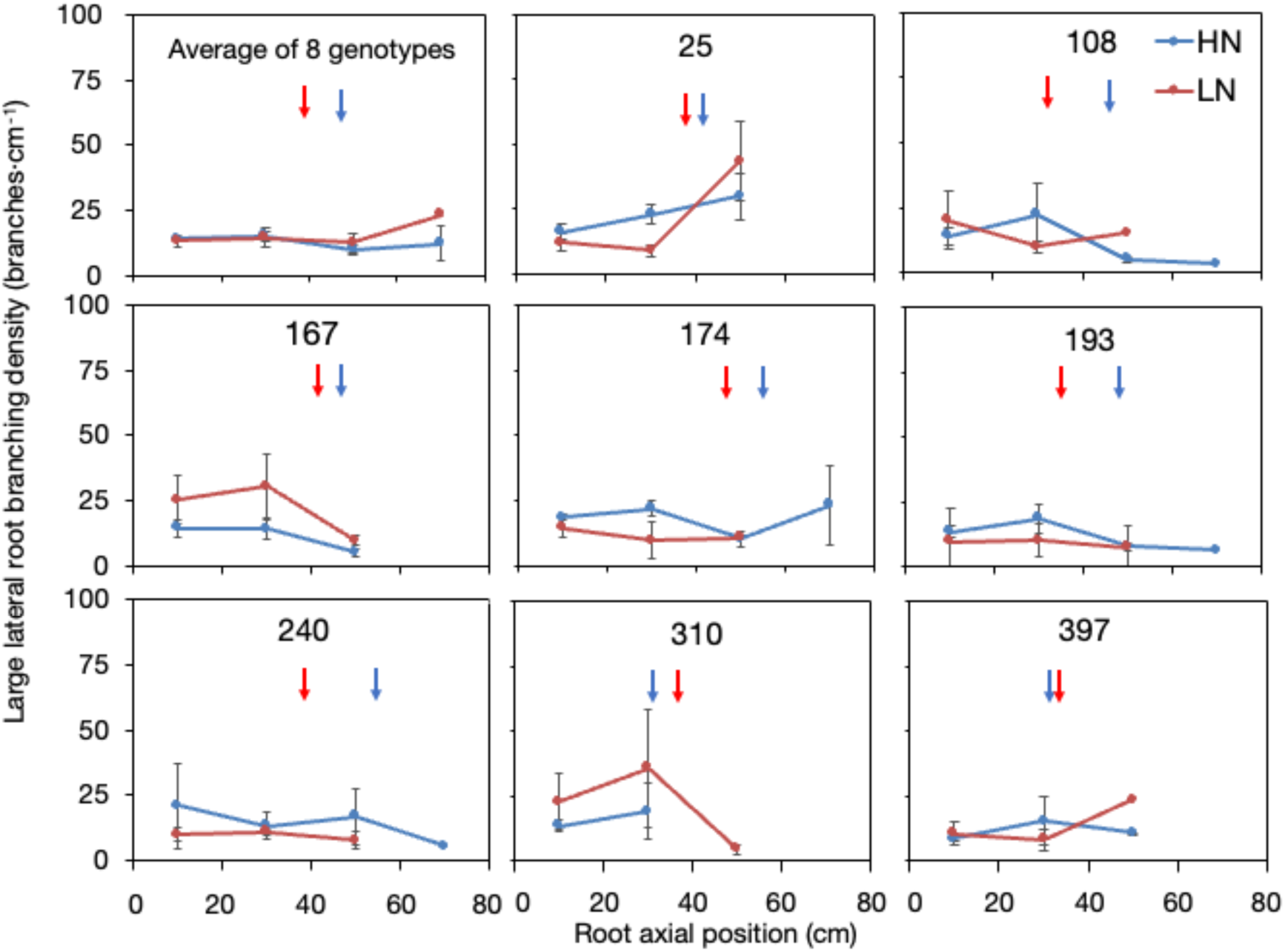
Effect of nitrogen treatment on distribution of large lateral root branching density along nodal root axes of each genotype in the RDP1 experiment. Values shown are means ± SE. Where no error bar is shown, there was only one nodal root sample at that depth. Arrows indicate mean nodal root length for high N (blue) and low N (red) treatments.

**Table 2.** The effects of genotype and root axial position on lateral root distribution of RDP1 lines.

|  | Large lateral root length (cm) |  | Large lateral root branching density |  | Small lateral root length (cm) |  |
| --- | --- | --- | --- | --- | --- | --- |
|  | F | P | F | P | F | P |
| Treatment (N) | 25.748 | <0.001 | 0.024 | 0.877 | 16.635 | <0.001 |
| Genotype (G) | 5.296 | <0.001 | 1.015 | 0.425 | 4.164 | <0.001 |
| Root axial position (RP) | 16.428 | <0.001 | 6.743 | <0.001 | 15.92 | <0.001 |
| N*G | 3.413 | 0.002 | 2.498 | 0.02 | 2.949 | 0.007 |
| N*RP | 4.161 | 0.018 | 0.065 | 0.937 | 4.859 | 0.01 |
| G*RP | 1.12 | 0.345 | 1.139 | 0.328 | 0.745 | 0.75 |
| N*G*RP | 0.973 | 0.483 | 2.127 | 0.018 | 0.572 | 0.871 |
| Root axial position | Mean | SE | Mean | SE | Mean | SE |
| 0-20 cm | 228.16 a | 20.13 | 16.57 b | 7.42 | 153.84 a | 13.32 |
| 20-40 cm | 217.25 a | 20.69 | 22.02 ab | 7.63 | 104.22 ab | 13.69 |
| 40-60 cm | 43.52 b | 28.14 | 68.23 a | 10.12 | 18.85 bc | 18.62 |
| 60-80 cm | 12.00 b | 71.68 | 12.17 b | 24.19 | 7.80 c | 47.44 |
|  | Small lateral root branching density |  | Proportion of large lateral root length to total root length |  | Proportion of small lateral root length to total root length |  |
|  | F | P | F | P | F | P |
| Treatment (N) | 0.66 | 0.418 | 0.008 | 0.93 | 0.661 | 0.418 |
| Genotype (G) | 0.508 | 0.827 | 0.015 | 1 | 0.093 | 0.999 |
| Root axial position (RP) | 0.138 | 0.937 | 55.315 | <0.001 | 6.112 | 0.001 |
| N*G | 2.123 | 0.047 | 0.019 | 1 | 0.46 | 0.861 |
| N*RP | 0.407 | 0.666 | 0.738 | 0.481 | 0.321 | 0.726 |
| G*RP | 0.596 | 0.888 | 0.965 | 0.502 | 0.464 | 0.964 |
| N*G*RP | 0.843 | 0.614 | 3.629 | <0.001 | 0.389 | 0.958 |
| Root axial position | Mean | SE | Mean | SE | Mean | SE |
| 0-20 cm | 63.35 a | 12.32 | 0.59 a | 0.03 | 0.77 a | 0.12 |
| 20-40 cm | 63.73 a | 12.67 | 0.38 b | 0.03 | 0.35 ab | 0.12 |
| 40-60 cm | 72.87 a | 16.8 | 0.05 c | 0.03 | 0.04 ab | 0.15 |
| 60-80 cm | 22.28 a | 40.18 | 0.01 c | 0.08 | 0.02 b | 0.38 |

Analysis of variance (ANOVA), means and standard errors (SE) are shown for lateral root distribution traits at different root axial positions: 0-20 cm, 20-40 cm, 40-60 cm and 60-80 cm. Different letters indicate significant differences at $p < 0.05$ . Proportions of small and large lateral roots are based on total length of all root classes per nodal root.

Small lateral root length varied significantly with axial position but not genotype or treatment in the RIL experiment (Table S4), while in the RDP1 experiment, there were significant interactions of N treatment with genotype and root sampling position (Table 2, Figure 6). Genotypes that had the greatest small lateral root length under high N had the greatest reductions with low N, especially in the upper segments. The only significant factors affecting small lateral root branching density were axial position in the RILs and nitrogen treatment X genotype in the RDP1 lines (Table 2, Table S4).

**Figure 6.**
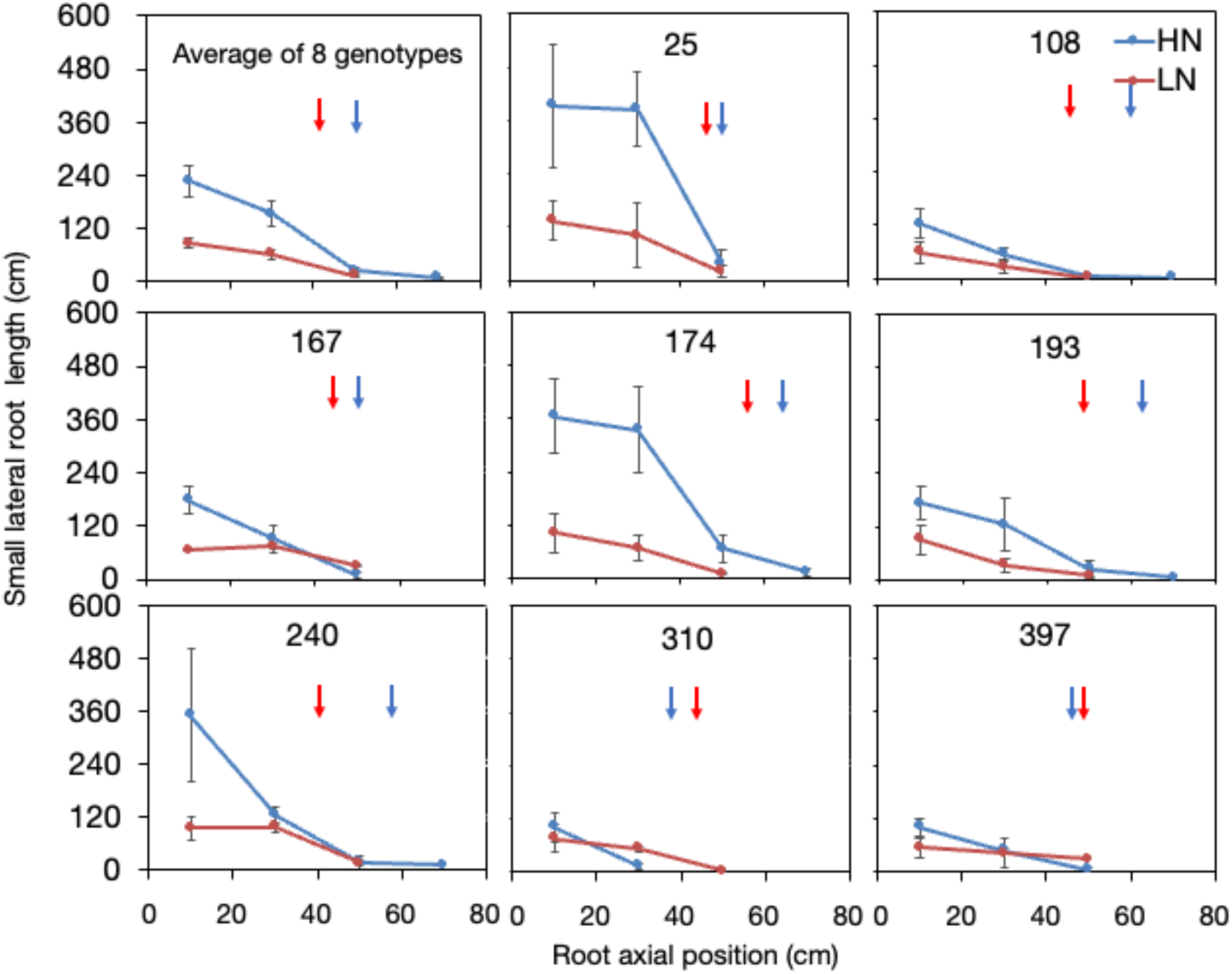
Effect of nitrogen treatment on distribution of small lateral root length along nodal root axes of each genotype in the RDP1 experiment. Values shown are means ± SE. Where no error bar is shown, there was only one nodal root sample at that depth. Arrows indicate mean nodal root length for high N (blue) and low N (red) treatments.

The proportion of large lateral root length to total root length per nodal root was significantly affected by root axial position, N treatment × root axial position and genotype × root axial position interactions in the RILs (Table S4), and there was a 3-way interaction in RDP1 (Table 2). Proportions of small lateral root length were only affected by sampling position.

### Root anatomical responses to low N

Anatomical responses of nodal roots to low nitrogen were assessed in the RIL experiment. For thick nodal roots, low N reduced RCSA, TSA, and MXV, and significantly interacted with genotype in its effects on %AA (Table 3). The strongest effects were on TSA, where genotype, treatment, and their interaction were all significant. Tissue was sampled at multiple distances from the base of thin nodal roots. There were significant nitrogen, genotype, and sampling position effects for most anatomy variables (Table 4). Significant treatment x genotype interactions were observed only for aerenchyma (AA and AA% (P<0.05), but differences among genotypes were not dramatic, with mean AA ranging from 0.08-0.17 mm^2^ and %AA ranging from 54-67% in the most mature tissue (20 cm sample, data not shown). As expected, aerenchyma was less developed in less mature root segments (Table 4).

**Table 3.** The effects of nitrogen treatment and genotype on anatomical phenotypes of thick nodal roots in the RIL experiment.

|  | RCSA (mm<sup>2</sup>) |  | AA (mm<sup>2</sup>) |  | AA% |  |
| --- | --- | --- | --- | --- | --- | --- |
|  | F | P | F | P | F | P |
| Treatment (N) | 6.270 | 0.016 | 1.313 | 0.258 | 0.247 | 0.622 |
| Genotype (G) | 1.344 | 0.251 | 1.363 | 0.243 | 1.913 | 0.089 |
| N*G | 1.827 | 0.104 | 1.864 | 0.097 | 3.019 | 0.011 |
| Treatments | Mean | SE | Mean | SE | Mean | SE |
| HN | 0.586 a | 0.03349 | 0.207 a | 0.019 | 32.53 a | 3.17 |
| LN | 0.489 b | 0.02336 | 0.181 a | 0.014 | 38.37 a | 2.84 |

|  | TSA (mm<sup>2</sup>) |  | MXA (mm<sup>2</sup> × 10<sup>-4</sup>) |  | MXV |  |
| --- | --- | --- | --- | --- | --- | --- |
|  | F | P | F | P | F | P |
| Treatment (N) | 17.051 | <0.001 | 3.830 | 0.056 | 18.947 | <0.001 |
| Genotype (G) | 4.558 | 0.001 | 1.707 | 0.130 | 1.579 | 0.065 |
| N*G | 2.610 | 0.023 | 1.029 | 0.424 | 1.952 | 0.082 |
| Treatments | Mean | SE | Mean | SE | Mean | SE |
| HN | 0.047 a | 0.003 | 17.03 a | 0.98 | 4.969 a | 0.131 |
| LN | 0.035 b | 0.002 | 14.73 a | 0.72 | 4.042 b | 0.188 |

Analysis of variance (ANOVA), means and standard errors (SE) are shown for root cross sectional area (RCSA), aerenchyma area (AA), aerenchyma area as percent of cortex (AA%), total stele area (TSA), mean metaxylem vessel area (MXA) and metaxylem vessel number (MXV). Different letters indicate significant difference at $p < 0.05$ .

**Table 4.** The effects of nitrogen treatment (N), genotype (G) and root axial position (RP) on anatomical phenotypes of thin nodal roots in the RIL experiment.

| Thin | RCSA (mm<sup>2</sup>) |  | AA (mm<sup>2</sup>) |  | AA% |  |
| --- | --- | --- | --- | --- | --- | --- |
|  | F | P | F | P | F | P |
| N Treatment (N) | 2.190 | 0.146 | 3.253 | 0.078 | 4.058 | 0.050 |
| Genotype (G) | 6.070 | <0.001 | 4.603 | 0.001 | 4.545 | 0.001 |
| Root axial position (RP) | 1.343 | 0.271 | 7.319 | 0.002 | 16.695 | <0.001 |
| N*G | 2.178 | 0.054 | 2.219 | 0.050 | 2.610 | 0.023 |
| N*RP | 0.446 | 0.643 | 0.372 | 0.692 | 0.074 | 0.928 |
| G*RP | 0.689 | 0.773 | 0.998 | 0.471 | 0.947 | 0.519 |
| Treatment | Mean | SE | Mean | SE | Mean | SE |
| HN | 0.277 a | 0.022 | 0.168 a | 0.015 | 59.92 a | 1.90 |
| LN | 0.221 b | 0.017 | 0.131 b | 0.011 | 56.81 a | 2.33 |
| Root axial position | Mean | SE | Mean | SE | Mean | SE |
| 20 cm | 0.222 a | 0.015 | 0.140 a | 0.011 | 60.715 a | 1.954 |
| 40 cm | 0.227 a | 0.018 | 0.134 a | 0.012 | 57.182 a | 2.342 |
| 60 cm | 0.196 a | 0.014 | 0.092 b | 0.008 | 45.201 b | 2.504 |
|  | TSA (mm<sup>2</sup>) |  | MXA (mm<sup>2</sup> × 10<sup>4</sup>) |  | MXV |  |
|  | F | P | F | P | F | P |
| N Treatment (N) | 19.314 | <0.001 | 5.927 | 0.019 | 18.400 | <0.001 |
| Genotype (G) | 14.357 | <0.001 | 8.266 | <0.001 | 4.238 | 0.001 |
| Root axial position (RP) | 8.009 | 0.001 | 6.746 | 0.003 | 9.898 | <0.001 |
| N*G | 2.135 | 0.058 | 1.942 | 0.084 | 0.733 | 0.645 |
| N*RP | 0.349 | 0.708 | 0.609 | 0.548 | 1.270 | 0.290 |
| G*RP | 1.552 | 0.131 | 2.069 | 0.033 | 0.806 | 0.658 |
| Treatment | Mean | SE | Mean | SE | Mean | SE |
| HN | 0.024 a | 0.002 | 13.690 a | 0.870 | 3.125 a | 0.209 |
| LN | 0.017 b | 0.001 | 11.940 a | 0.950 | 2.229 b | 0.184 |
| Root axial position | Mean | SE | Mean | SE | Mean | SE |
| 20 cm | 0.022 a | 0.002 | 11.12 ab | 0.55 | 2.969 a | 0.193 |
| 40 cm | 0.020 a | 0.002 | 12.68 a | 0.85 | 2.323 b | 0.219 |
| 60 cm | 0.016 b | 0.001 | 9.97 b | 0.78 | 1.936 b | 0.167 |

Analysis of variance (ANOVA), means and standard errors (SE) are shown for anatomical traits for thin nodal roots. Different letters indicate significant difference ( $p < 0.05$ ). None of the three-way interactions (N\*G\*RP) were significant.

Mean values for TSA and MXV were significantly reduced under low nitrogen, and there was a significant genotype × sampling position interaction for TSA (Table 4). There were no significant effects of genotype, N treatment, or sampling position on anatomical traits of large lateral roots (data not shown).

### Allometric analysis of root phenotypes

Root phenotypes were subject to allometric analyses to determine to what extent they were determined by overall plant size as opposed to specific responses to low nitrogen. Root architectural traits were hyperallometric (alpha>0.33) or not significant (Table 5). The strongest allometric relationship was between nodal root number and shoot biomass, which was strongly hyperallometric in both experiments and in both N treatments, indicating a disproportionately large investment in nodal root number relative to shoot growth. These values were greater with low N, indicating that low N stress accelerated the development of nodal roots relative to shoot growth. Lateral root phenotypes showed no disproportionate responses in the RILs experiment, and hyperallometric responses with low nitrogen in the RDP1 experiment, though these were less pronounced than the nodal root responses (Table 5).

**Table 5.** Allometric partitioning coefficients for root phenotypes vs. shoot biomass in the RILs and RDP1 experiments. For traits with significant p-values, the response is considered hyperallometric if the alpha value is >0.33 and hypoallometric if the alpha value is <0.33.

| Experiment | Trait | High N |  |  |  | Low N |  |  |  |
| --- | --- | --- | --- | --- | --- | --- | --- | --- | --- |
| | | Adj. R<sup>2</sup> | alpha | $p$-value | n | Adj. R<sup>2</sup> | alpha | $p$-value | n |
| RILs | Nodal root number | 0.5627 | 1.0336 | $<0.001$ | 32 | 0.4359 | 1.1389 | $<0.001$ | 32 |
|  | Thin nodal root length | 0.13 | 0.8386 | 0.046 | 32 | 0.277 | 0.1218 | 0.001 | 32 |
|  | Thick nodal root length | 0.0221 | 0.1201 | 0.135 | 32 | 0.2379 | 0.6662 | 0.002 | 32 |
|  | Large lateral root length per thin nodal root | 0.0159 | 0.0174 | 0.385 | 32 | 0.0686 | -0.2176 | 0.074 | 32 |
|  | Large lateral root length per thick nodal root | 0.0591 | 0.0982 | 0.226 | 24 | 0.1739 | 0.194 | 0.432 | 16 |
|  | Small lateral root length per thin nodal root | 0.0778 | 0.1152 | 0.206 | 32 | 0.0063 | -0.0399 | 0.334 | 32 |
|  | Small lateral root length per thick nodal root | 0.0336 | 0.0564 | 0.126 | 32 | 0.0429 | -0.0779 | 0.127 | 32 |
|  | Thin nodal root cross-sectional area | 0.021 | 0.0685 | 0.45 | 32 | 0.0014 | 0.0243 | 0.422 | 31 |
| RDP1 | Nodal root number | 0.3422 | 0.8092 | 0.001 | 27 | 0.7133 | 1.5806 | $<0.001$ | 22 |
| | Nodal root length | 0.3399 | 1.1084 | 0.001 | 28 | 0.4048 | 1.7441 | $<0.001$ | 25 |
| | Large lateral root length | 0.4415 | 0.3646 | $<0.001$ | 28 | 0.4668 | 0.6369 | $<0.001$ | 25 |
| | Small lateral root length | 0.4896 | 0.365 | $<0.001$ | 28 | 0.4194 | 0.5746 | $<0.001$ | 25 |

## Discussion

Rice roots showed significant genotypic variation in root phenotypes and in plasticity of root phenotypes in response to low nitrogen. Many of these changes were correlated with biomass reduction, i.e. they were isometric. In this paper, we focus on phenotypes that exhibited genotype X nitrogen treatment interactions, since these could be of use to breeders targeting improved nitrogen efficiency.

Lynch has proposed a *steep, cheap and deep* ideotype for efficient acquisition of nitrogen and water by maize crops (Lynch 2013), several components of which have proven beneficial for maize performance in soils with low nitrogen fertility (Saengwilai et al. 2014b; Zhan and Lynch 2015; York and Lynch 2015; Lynch et al. 2023). We hypothesized that under nitrogen leaching conditions, rice plants, like maize plants, would also benefit from root traits that allow deeper soil exploration, e.g. allocation of resources to root elongation vs. number of axial and lateral roots and reducing the metabolic cost of soil exploration via anatomical phenotypes such as cortical aerenchyma, reduced cortical cell file number, and greater cell size. Beneficial phenotypes could be stable or plastic, *i.e.* they could change in response to nitrogen stress. In our experiments, rice was grown in mesocosms that simulated the nitrogen leaching that occurs in field conditions, so that low nitrogen mesocosms had less nitrogen overall, but nitrogen became increasingly more depleted in the shallower part of the mesocosms than in the deeper soil as plant growth progressed. We therefore expected that genotypes that were able to effectively capture nitrogen in the deeper parts of the mesocosms would perform better under low nitrogen.

Nitrogen efficiency could be improved by greater uptake of nitrogen per unit surface area of root via anatomical and molecular mechanisms, from greater investment in root classes or segments (e.g. increased branching resulting in greater proportion of young root axes) that have greater uptake rates, or from placement of roots in soil domains with greater nitrogen availability (Griffiths and York 2020). All of these interacting mechanisms could be operational in rice. Nitrogen content per unit root length or root dry weight was about four-fold greater in low nitrogen (Table S3), which could indicate greater uptake efficiency or less transport of nitrogen to the shoot, perhaps to support increased investment in root development. We did not observe genotypic variation or interactions of genotype and nitrogen treatment for these metrics, however, indicating that genotypic variation in biomass accumulation under low nitrogen could not be explained by changes in uptake over the entire root system. It is therefore more likely that root architecture and anatomy account for variation in performance under low nitrogen among genotypes. Rice root systems became relatively larger (greater root to shoot ratio) under low nitrogen, suggesting that greater investment in root development was an overall strategy, but again, there was no significant genetic variation for low nitrogen response (no N X G interaction). Therefore, more specific root changes to root architecture or anatomy should be important.

One aspect of the *steep, cheap and deep* ideotype is that fewer nodal roots should be beneficial under low nitrogen, improving deep soil exploration by prioritizing axial elongation over growth of additional nodal roots (Lynch 2013). In maize, genotypes with fewer crown (subterranean nodal) roots had a greater proportion of root length in deeper soil domains, better nitrogen acquisition, more biomass and greater yield under moderate nitrogen stress (Saengwilai et al. 2014b). In our experiments, nodal root number was positively correlated with shoot biomass under both high and low nitrogen (r = 0.58-0.76). Nodal root number and nodal root length were not correlated in high nitrogen and were positively correlated in low nitrogen (r=0.44 in RILs, r=0.55 in RDP1). Therefore, in most rice genotypes, nodal root number was proportional to overall plant growth, and fewer nodal roots was not associated with an increase in axial root length or deep soil exploration. An exception was RDP1 line 174 (Azucena), which had the greatest biomass and the fewest nodal roots of the RDP1 lines under low nitrogen (Figures 1-2).

An important difference between rice and maize is the presence of tillering in rice. Each tiller develops nodal roots synchronized with leaf formation (Rebouillat et al. 2009), so that tillering and nodal root production are correlated. In our studies, these correlations were similar across nitrogen treatments and populations (r values from 0.61-0.67).

Although phenotypes with fewer nodal roots were not generally advantageous for nitrogen acquisition in rice, nodal root length and root depth should be associated with dry matter accumulation, particularly as nitrogen was depleted from the upper soil during the later stages of growth. Nodal root phenotypes were measured in detail in the RILs experiment (Table 1). Thin nodal roots were about three times longer than thick nodal roots and bore much greater length of large and small lateral roots. Thin nodal roots were therefore much more likely to contribute to nitrogen uptake throughout the soil profile, both directly and via their lateral roots. Nodal root lengths were reduced by low nitrogen, but to a much lesser extent than reductions in shoot biomass. Both thin and thick nodal root lengths were positively correlated with biomass accumulation over all genotypes (r=0.48-0.52). These longer nodal roots were likely to be important for nitrogen acquisition in the deeper strata of the mesocosms, where far fewer roots were found. These results support previous *OpenSimRoot/Rice* simulations of root architectural phenotypes associated with rice growth under low nitrogen. In simulations, all phenotypic clusters that had superior performance to the reference genotype, IR64, had the same or greater numbers of nodal roots than the reference phenotype, IR64 (Ajmera et al. 2022). A second aspect of the *steep, cheap and deep* ideotype is that fewer, longer lateral roots should be associated better N capture than many, shorter lateral roots. *SimRoot* modeling indicated that compared with optimal density for abundant nitrogen availability, optimal lateral root branching density for maize grown with moderate nitrogen stress was slightly less and with severe nitrogen stress much less (Postma et al. 2014). These results were confirmed in greenhouse and field experiments, where maize genotypes with few, long lateral roots showed superior performance under low nitrogen to those with many, short lateral roots (Zhan and Lynch 2015). However, simulation of rice growth with low nitrogen predicted that phenotypes with denser large lateral roots would perform better under both low and intermediate N supply, while small lateral root density had little impact (Ajmera et al. 2022). In both of our experiments, there was significant genetic variation in large lateral root density and length, and both traits were associated with better performance under low nitrogen (r=-0.43-0.61). There was significant genotypic variation in large lateral root responses to low nitrogen in the RDP1 experiment but not in the RILs experiment (Table 1, Table S2). Genotypes that maintain root elongation with low nitrogen should have improved access to the N remaining in deep soil. In rice, which has more abundant nodal roots, both nodal roots and large lateral roots could contribute to deep soil exploration, and much of the nodal root contribution could be via the subtending large lateral roots. For both types of axial roots, favoring development of subtending lateral roots on the deeper root segments would be particularly advantageous in capturing nitrogen from deeper soil strata.

The “cheap” aspect of the *steep, cheap and deep* ideotype posits that phenes contributing to reduced metabolic cost per unit of root length should allow for greater soil exploration. One such trait is aerenchyma formation, which reduces metabolic cost by lysis of cortical cells. Aerenchyma formation is associated with reduced respiration rates per unit root length in maize (Fan et al. 2003; Jaramillo et al. 2013; Saengwilai et al. 2014a) and rice (Fonta et al. 2022a). Previous research with a single rice cultivar grown in nutrient solution showed that nitrogen starvation reduced aerenchyma formation (Abiko and Obara 2014). In our study in an aerobic solid medium, we observed genetic variation for both the extent and nitrogen responsiveness of aerenchyma formation (Table 3). Aerenchyma formation may have contributed to performance differences. Another factor affecting root metabolic cost is root diameter, captured in the RILs experiment as nodal root cross-sectional area (RCSA) and in the RDP1 experiment as root diameter (from WinRhizo). Rice nodal root diameter was previously shown to be reduced by water deficit (Kadam et al. 2017) and low phosphorus stress (Vejchasarn et al. 2016). Although RCSA was less in low nitrogen treatments in the RILs experiment, it was unaffected in the RDP1 experiment, and there was no significant interaction between genotype and nitrogen treatment on RCSA or root diameter, indicating that this was unlikely to be a factor affecting genetic differences in performance.

Root phenotypes that are beneficial with nitrogen stress in leaching soils should also be useful with drought stress in terminal drought environments. In both cases, a limiting resource becomes increasingly depleted over time in the upper soil strata, due to plant uptake and nitrogen leaching with rainfall, and plant uptake and evaporation of moisture from the surface during drought. Several studies demonstrate that root phenes leading to deeper roots improve maize performance under drought (Chimungu et al. 2014a, b; Zhan et al. 2015; Gao and Lynch 2016; Klein et al. 2020). In rice, deep rooting is well established as a desirable phenotype during drought, especially in upland systems (Gowda et al. 2011; Uga et al. 2015). Deep rooting in rice has been associated with steeper angles of nodal roots (Kato et al. 2006; Uga et al. 2013), maintenance or increase of allometric scaling between nodal root number and shoot biomass, and increased root length density in deeper soil (Kijoji et al. 2013; Kameoka et al. 2015). Recent work with rice and maize has demonstrated that integrated phenotypes, i.e. specific combinations of traits, are useful to explain variation in performance with drought stress (Klein et al. 2020; Fonta et al. 2022b). Although this study did not include enough genotypes for a cluster analysis of integrated phenotypes, these results and those of (Fonta et al. 2022b) suggest that deep rooting is facilitated by maintaining elongation of nodal roots and large lateral roots, and development of cost-saving anatomical features such as aerenchyma.

### Conclusion

Rice lines demonstrated significant genotypic variation for root responses to low nitrogen stress. Stratified nitrogen availability in overall nitrogen-limiting conditions substantially reduced overall biomass accumulation, but there was genetic variation for the biomass impact of low nitrogen as well as the responses of several architectural and anatomical traits. Under nitrogen leaching conditions, responses resulting in greater development of deep root length and reduced root cost could be beneficial for crop performance.

## Notes

### Competing Interest Statement

The authors have declared no competing interest.

