## Supplemental figure and tables for "Responses of root architectural and anatomical traits to low nitrogen stress in rice"

### Supplemental Information Tian, Lynch, Brown

**Supplemental Figure 1** Soil nitrate levels over time at various depths within mesocosms supplied with high (HN) (a) or low (LN) nitrogen (b) for RDP1, and HN (c) or LN (d) for RILs, with 4 replications.

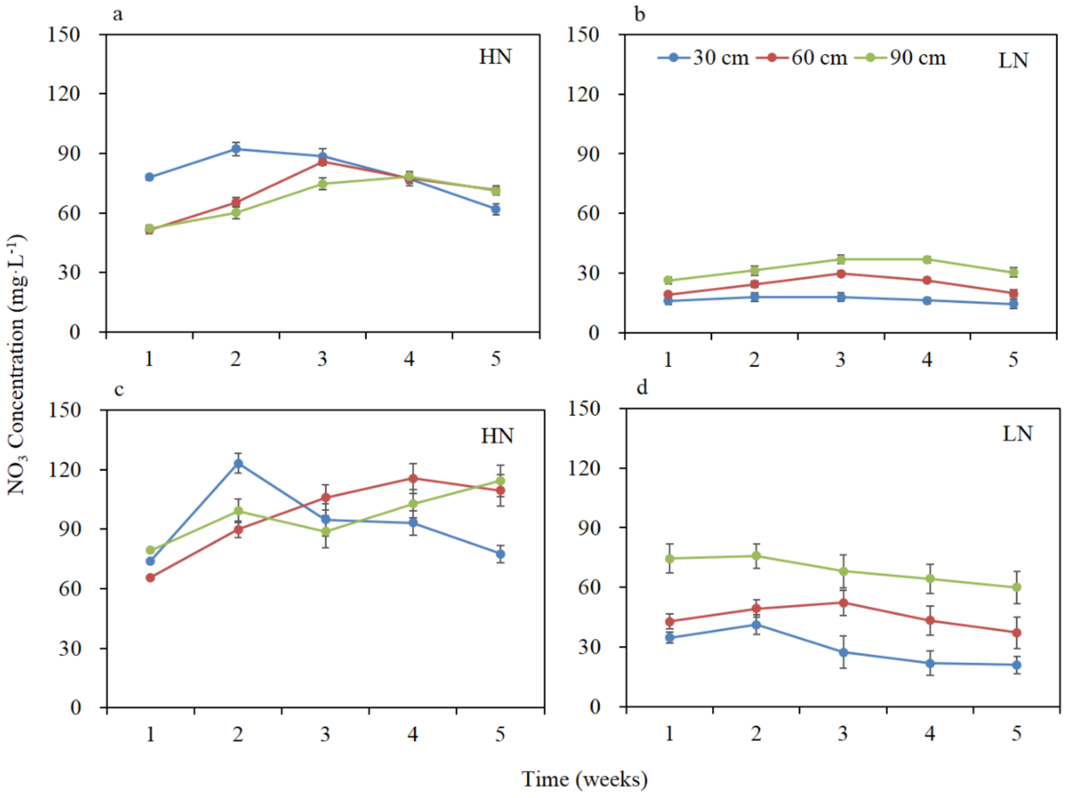

**Table S1** Rice cultivars used for RDP1 experiment

| Number | Cultivar Name | Origin | GSOR ID |
| --- | --- | --- | --- |
| 25 | Carolina Gold | United States | GSOR#301023 |
| 108 | Moroberekan | Guinea | GSOR#301100 |
| 167 | B6616A4-22-Bk-5-4 | United States | GSOR#301158 |
| 174 | Azucena | Philippines | GSOR#301165 |
| 193 | Fossa Av | Burkina Faso | GSOR#301184 |
| 240 | WAB 501-11-5-1 | Cote D'Ivoire | GSOR#301230 |
| 310 | R 101 | Zaire | GSOR#301300 |
| 397 | Cybonnet | United States | GSOR#301380 |

**Table S2** The effects of nitrogen treatment and genotype on shoot and root morphological phenotypes of RDP1 lines.

| | Shoot biomass (g·plant$^{-1}$) | | Number of leaves (plant$^{-1}$) | | Number of tillers (plant$^{-1}$) | | SPAD | |
|---|---|---|---|---|---|---|---|---|
| | F | P | F | P | F | P | F | P |
| Treatment (N) | 367.863 | <0.001 | 177.692 | <0.001 | 200.653 | <0.001 | 180.438 | <0.001 |
| Genotype (G) | 69.014 | <0.001 | 3.654 | 0.004 | 6.429 | <0.001 | 1.641 | 0.150 |
| N*G | 39.245 | <0.001 | 2.597 | 0.026 | 3.151 | 0.009 | 1.353 | 0.250 |
| Treatments | Mean | SE | Mean | SE | Mean | SE | Mean | SE |
| HN | 1.204 a | 0.082 | 16.2 a | 0.81 | 4.8 a | 0.29 | 42.02 a | 0.43 |
| LN | 0.218 b | 0.025 | 6.3 b | 0.29 | 1.5 b | 0.14 | 30.40 b | 0.78 |

| | Root dry weight (g·plant$^{-1}$) | | Root to shoot ratio | | Maximum root length (cm) | | Nodal root number (plant$^{-1}$) | |
|---|---|---|---|---|---|---|---|---|
| | F | P | F | P | F | P | F | P |
| Treatment (N) | 50.779 | <0.001 | 15.88 | <0.001 | 12.339 | 0.001 | 78.326 | <0.001 |
| Genotype (G) | 3.585 | 0.004 | 0.494 | 0.830 | 5.146 | <0.001 | 2.801 | 0.018 |
| N*G | 1.709 | 0.135 | 0.070 | 0.999 | 2.591 | 0.027 | 2.182 | 0.057 |
| Treatments | Mean | SE | Mean | SE | Mean | SE | Mean | SE |
| HN | 0.367 a | 0.024 | 0.26 b | 0.04 | 59.1 a | 15.260 | 31.03 a | 1.64 |
| LN | 0.114 b | 0.026 | 0.52 a | 0.05 | 47.7 b | 16.766 | 15.69 b | 0.87 |

| | Nodal root length (cm) | | Small lateral root length (cm) | | Large lateral root length (cm) | | Root diameter (mm) | |
|---|---|---|---|---|---|---|---|---|
| | F | P | F | P | F | P | F | P |
| Treatment (N) | 24.044 | <0.001 | 41.476 | <0.001 | 46.311 | <0.001 | 1.607 | 0.213 |
| Genotype (G) | 4.210 | 0.002 | 9.659 | <0.001 | 9.293 | <0.001 | 2.402 | 0.039 |
| N*G | 1.658 | 0.151 | 5.069 | 0.001 | 5.721 | <0.001 | 0.925 | 0.499 |
| Treatments | Mean | SE | Mean | SE | Mean | SE | Mean | SE |
| HN | 50.0 a | 2.11 | 439.7 a | 69.17 | 685.0 a | 99.61 | 0.30 a | 0.01 |
| LN | 39.0 b | 1.78 | 162.1 b | 25.20 | 261.5 b | 30.56 | 0.32 a | 0.02 |

Analysis of variance (ANOVA), means and standard errors (SE) are shown. Different letters indicate significant differences at $p<0.05$. Lateral root traits are for thin nodal roots.

**Table S3** The effects of nitrogen treatment and genotype on N content of RILs

| | Shoot N content (mg·plant$^{-1}$) | | Root N content (mg·plant$^{-1}$) | | Shoot N content per unit root length (mg·m$^{-1}$) | | Shoot N content per unit root dry weight (mg·g$^{-1}$) | |
|---|---|---|---|---|---|---|---|---|
| | F | P | F | P | F | P | F | P |
| Treatment (N) | 329.952 | <0.001 | 120.475 | <0.001 | 19.291 | <0.001 | 25.209 | <0.001 |
| Genotype (G) | 2.525 | 0.027 | 4.132 | 0.001 | 0.608 | 0.746 | 0.442 | 0.871 |
| N*G | 1.743 | 0.121 | 1.854 | 0.098 | 0.352 | 0.925 | 0.294 | 0.953 |
| Treatments | Mean | SE | Mean | SE | Mean | SE | Mean | SE |
| HN | 77.92 a | 3.77 | 16.85 a | 1.25 | 0.116 b | 0.013 | 45.67 b | 6.28 |
| LN | 13.39 b | 1.29 | 4.37 b | 0.56 | 0.476 a | 0.076 | 184.06 a | 24.74 |

N content is calculated as a fraction of total root system length or weight.

**Table S4** The effects of nitrogen treatment, genotype and root axial position on lateral root distribution in the RIL experiment.

| | Small lateral root length (cm) | | Large lateral root length (cm) | | Small lateral root branching density | |
|---|---|---|---|---|---|---|
| | F | P | F | P | F | P |
| Treatment (N) | 0.143 | 0.706 | 0.714 | 0.400 | 0.039 | 0.843 |
| Genotype (G) | 0.870 | 0.532 | 1.706 | 0.112 | 0.374 | 0.916 |
| Root axial position (RP) | 14.208 | <0.001 | 18.521 | <0.001 | 5.586 | 0.001 |
| N*G | 1.123 | 0.352 | 0.873 | 0.530 | 1.814 | 0.089 |
| N*RP | 0.304 | 0.822 | 0.720 | 0.541 | 0.478 | 0.698 |
| G*RP | 0.712 | 0.801 | 0.878 | 0.610 | 0.379 | 0.992 |
| N*G*RP | 0.364 | 0.992 | 0.615 | 0.883 | 0.424 | 0.981 |
| Root axial position | Mean | SE | Mean | SE | Mean | SE |
| 0-20 cm | 176.21 a | 19.93 | 232.11 a | 22.96 | 185.94 ab | 19.77 |
| 20-40 cm | 191.87 a | 17.94 | 294.84 a | 18.12 | 234.74 a | 34.32 |
| 40-60 cm | 63.17 b | 12.52 | 140.23 b | 18.69 | 104.72 bc | 18.65 |
| 60-80 cm | 11.93 b | 4.61 | 30.55 c | 12.83 | 17.58 c | 6.09 |
| Treatments | Mean | SE | Mean | SE | Mean | SE |
| HN | 122.15 a | 13.98 | 196.17 a | 15.23 | 151.36 a | 20.47 |
| LN | 116.09 a | 15.50 | 176.77 a | 17.32 | 149.19 a | 23.28 |

| | Large lateral root branching density | | Proportion of small lateral root length to total root length | | Proportion of large lateral root length to total root length | |
|---|---|---|---|---|---|---|
| | F | P | F | P | F | P |
| Treatment (N) | 0.019 | 0.889 | 0.149 | 0.700 | 0.558 | 0.457 |
| Genotype (G) | 1.028 | 0.414 | 0.809 | 0.581 | 0.475 | 0.851 |
| Root axial position (RP) | 6.693 | <0.001 | 29.430 | <0.001 | 48.465 | <0.001 |
| N*G | 0.696 | 0.675 | 0.496 | 0.836 | 0.819 | 0.573 |
| N*RP | 0.448 | 0.719 | 0.726 | 0.538 | 3.017 | 0.032 |
| G*RP | 0.453 | 0.976 | 1.164 | 0.297 | 1.708 | 0.042 |
| N*G*RP | 0.573 | 0.914 | 0.296 | 0.998 | 0.598 | 0.896 |
| Root axial position | Mean | SE | Mean | SE | Mean | SE |
| 0-20 cm | 19.62 a | 1.36 | 16.37 a | 1.52 | 21.62 ab | 1.56 |
| 20-40 cm | 24.65 a | 2.88 | 17.05 a | 1.06 | 27.61 a | 1.10 |
| 40-60 cm | 16.26 a | 2.04 | 4.85 b | 0.61 | 11.05 c | 1.14 |

| | | | | | | |
|---|---|---|---|---|---|---|
| 60-80 cm | 5.92 b | 1.97 | 0.54 b | 0.17 | 1.70 d | 0.59 |
| Treatment | Mean | SE | Mean | SE | Mean | SE |
| HN | 17.34 a | 1.73 | 10.10 a | 0.90 | 15.83 a | 0.94 |
| LN | 17.77 a | 1.97 | 10.87 a | 1.02 | 17.57 a | 1.07 |

Analysis of variance (ANOVA), means and standard errors (SE) are shown for lateral root distribution at different root positions (RP): 0-20 cm, 20-40 cm, 40-60 cm and 60-80 cm. Different letters indicate significant difference ($p$<0.05)
